# Buffer Therapy in Acute Metabolic Acidosis: Effects on Glomerular Permeability and Acid-Base Status

**DOI:** 10.1101/528521

**Authors:** Jan Schnapauff, David Piros, Anna Rippe, Peter Bentzer, Naomi Clyne, Carl M. Öberg

## Abstract

Correction of acute metabolic acidosis using sodium bicarbonate is effective, but has been hypothesized to exacerbate intra-cellular acidosis causing cellular dysfunction. The effects of acidemia and bicarbonate therapy on the cellular components of the glomerular filtration barrier, crucial for the integrity of the renal filter, are as yet unknown. Controversy persists regarding the most appropriate method to assess acid-base status: the “Stewart approach” or the “Siggaard-Andersen approach” using the standard base excess (SBE). Here we performed physiological studies in anesthetized Sprague-Dawley rats during severe metabolic acidosis (HCl iv 6 mmol kg^−1^) and following bicarbonate (2.5 mmol kg^−1^) administration. We assessed glomerular permeability using sieving coefficients of polydisperse fluorescein isothiocyanate (FITC)-Ficoll 70/400. Acid-base status was evaluated using SBE, standard bicarbonate, total CO_2_, the Stewart-Fencl strong ion difference (ΔSID = Na – Cl – 38) and a theoretical model of plasma and erythrocyte strong ion difference. Our data show that neither acidosis nor its correction with NaHCO_3_ altered glomerular permeability. We identified ΔSID as a strong estimator of plasma base excess (as assessed using the Van Slyke equation). *In silico* modeling indicates that changes in the strong ion difference in erythrocytes would explain their buffering effect by means of a shift of anions from the extracellular fluid. These data demonstrate a remarkable tolerance of the glomerular filter to severe acute acidosis and bicarbonate therapy. Our results also cast light on the buffer mechanism in erythrocytes and the ability of different acid-base parameters to evaluate the extent of an acid-base disorder.

## INTRODUCTION

Metabolic acidosis is associated with poor outcome in critically ill patients with reported mortality rates exceeding 50% when the pH stays below 7.20 (18). Bicarbonate therapy is a simple and effective treatment to correct acute acidemia and chronic buffer depletion. However, the safety and feasibility of buffer treatment in acute metabolic acidosis are to date little studied and remain controversial (23). Adverse effects of bicarbonate therapy are well-described in the literature and include hypokalemia (2), hypocalcemia (42), hypercapnia (1), hypernatremia and fluid overload (22). Buffer therapy will also, by mitigating acidemia, lead to a left-shift of the oxygen dissociation curve (reduced Bohr effect) with increased binding of oxygen to hemoglobin and reduced delivery to tissues, potentially contributing to an increase in tissue hypoxia. On the other hand, it is well-recognized that acute metabolic acidosis can have direct negative effects on a number of organ systems, especially in the form of reduced cardiac output and contractility, which mainly occurs at a pH < 7.10-7.20 (29).

The method by which the extent of a metabolic acid-base disorder should be determined was the topic of the historical “Great Trans-Atlantic Acid-base Debate” between the North American “Boston school” (favoring bicarbonate) and the European “Copenhagen school” (favoring base excess) which also comprise the buffer effects of erythrocyte hemoglobin (38). New controversy emerged by the introduction of a “Canadian school” with the so called “Stewart approach” described by the Canadian physiologist Stewart in the 1980ies (37). This method, which does not include erythrocyte buffers, is based on the balance between blood plasma buffer ions (usually conjugate base anions of weak acids such as carbonic acid) and *strong ions*, which do not act as buffers. Due to the complexity of the original Stewart model, a simplified methodology, the so called Fencl-Stewart approach has been proposed, introducing a simplified version of the SID = Na – Cl. The normal SID between strong ions in blood plasma is ~38 meq/L (25 meq/L bicarbonate, 12 meq/L albumin and 1 meq/L phosphate) and thus the strong ion gap ΔSID = Na – Cl – 38 is an approximation of the amount of buffer ions in plasma.

Metabolic acidosis is an invariable feature of acute and chronic kidney disease and results mainly from reduced ammoniagenesis and impaired tubular reabsorption of bicarbonate. The observation that a drop in plasma pH could cause proteinuria is not new, and there are some reports (15, 43) showing that metabolic acidosis leads to increased proteinuria. In an early study, Gardner (15) provided data that indicated alterations in protein transport at the glomerular level during acidemia. In contrast, Throssell and colleagues (43) found significant low molecular weight proteinuria in acidotic rats, implying a tubular origin. Recent data reveal that the integrity of the glomerular filter is crucially determined by the concerted actions of the cellular components of the glomerulus (4, 14, 40), especially podocytes (16), which appear to tightly regulate glomerular permeability. Moreover, several *in vitro* and *in vivo* studies have indicated that both extra-cellular acidemia and bicarbonate therapy can exacerbate intra-cellular acidosis (22, 24), conceivably leading to cellular dysfunction. Thus, we hypothesized that both acidemia and its correction using bicarbonate, could affect glomerular permeability. Also, recent data indicate that the buffering effect of erythrocytes is facilitated chiefly via the Anion exchanger 1 (AE1) (41), making up to 25% of the red cell membrane surface, with no involvement of other transporters. On this basis, we developed a strong ion theory that includes the buffering effect of erythrocytes and show that it can explain the difference between ΔSID and SBE in our data. We also investigated the capability of several parameters to assess buffer deficit, since a number of different parameters for acid-base status are currently in clinical use.

## METHODS

### Animals

Experimental studies were performed in 8 male Sprague-Dawley rats (Möllegard, Lille Stensved, Denmark) having an average body weight (BW) of 269 g (255-289 g) with free access to food and water. The local Animal Ethics Committee at Lund University, Sweden approved all procedures.

### Surgery

Induction of anesthesia was performed by means of an intraperitoneal injection with pentobarbital (Pentobarbitalnatrium vet. APL 60 mg mL^−1^), 90 mg kg^−1^ BW, and maintained by administration of pentobarbital (3-6 mg) via the tail artery as needed. A heating pad was used to maintain the body temperature at 37°C. A tracheotomy was performed (using a PE-240 tube) and connected to a ventilator (Ugo Basile; Biological Research Apparatus, Comerio, Italy) and ventilated in volume-controlled mode using a positive end-expiratory pressure of 4 cm water, respiratory rate 65-70 min^−1^ and a tidal volume of 3 mL kg^−1^. The tail artery was cannulated (using a PE-50 cannula) and used for administration of furosemide (Furosemid Recip, Årsta, Sweden), maintenance of anesthesia, and for continuous monitoring and registration of mean arterial blood pressure and heart rate (HR) on a data acquisition system (BioPack Systems model MP150 with AcqKnowledge software version 4.2.0, BioPack Systems Inc., Goleta, CA). The left carotid artery was cannulated (PE-50) and used to obtain blood samples for measurement of arterial blood gases and electrolyte concentrations. Hematocrit was determined by centrifugation of standard glass capillaries. The left and right internal jugular vein were cannulated (PE-50) for the administration of Ficoll and pharmacological interventions (HCl and sodium bicarbonate), respectively. A minimal abdominal incision (6-8 mm) was made to gain access to the left ureter. A very small dose of furosemide (0.375 mg kg^−1^) was given to facilitate cannulation and the ureter was then dissected free and cannulated using a PE-10 cannula, which was used for urine sampling. All connectors, cannulas and tubes in contact with the circulation were prepared and flushed post administration with a dilute heparin (Heparin LEO 5000 IE/mL) solution (~25 IE/mL in NaCl aq. 9 mg/mL) to prevent clotting.

### Experimental metabolic acidosis

All experiments commenced with a resting period of 20 min following surgery (Figure 1). The left carotid artery was cannulated and utilized for arterial blood samples for analysis of electrolytes, *p*CO_2_, urea, Hb, and pH using the I-STAT EC8+ cassette (I-STAT; Abbot Point of Care Inc, Abbot Park, IL). To prevent respiratory compensation of the metabolic acidosis, neuromuscular blockade was then induced with rocuronium bromide at a dose of 0.6 mg/kg (Esmeron, Merck Sharp & Dohme B.V., Haarlem, Netherlands) followed by a dose of 0.3 mg/kg every 40 min. The animal was carefully observed during the course of the experiment to ensure that no spontaneous breathing occurred. After an initial 5 min period for control (baseline) measurements, hydrochloric acid 6 mmol/kg (Saltsyra APL 1 mmol/mL, APL, Stockholm, Sweden) was given as an infusion during a period of 60 minutes starting directly after the baseline measurement period. After the infusion had stopped, new measurements were performed during a 5 min period after which sodium bicarbonate 2.5 mmol/kg (Natriumbikarbonat Fresenius Kabi 0.6 mmol/mL, Uppsala, Sweden) was given as an intravenous infusion during a period of 10 minutes.

**Figure 1.**
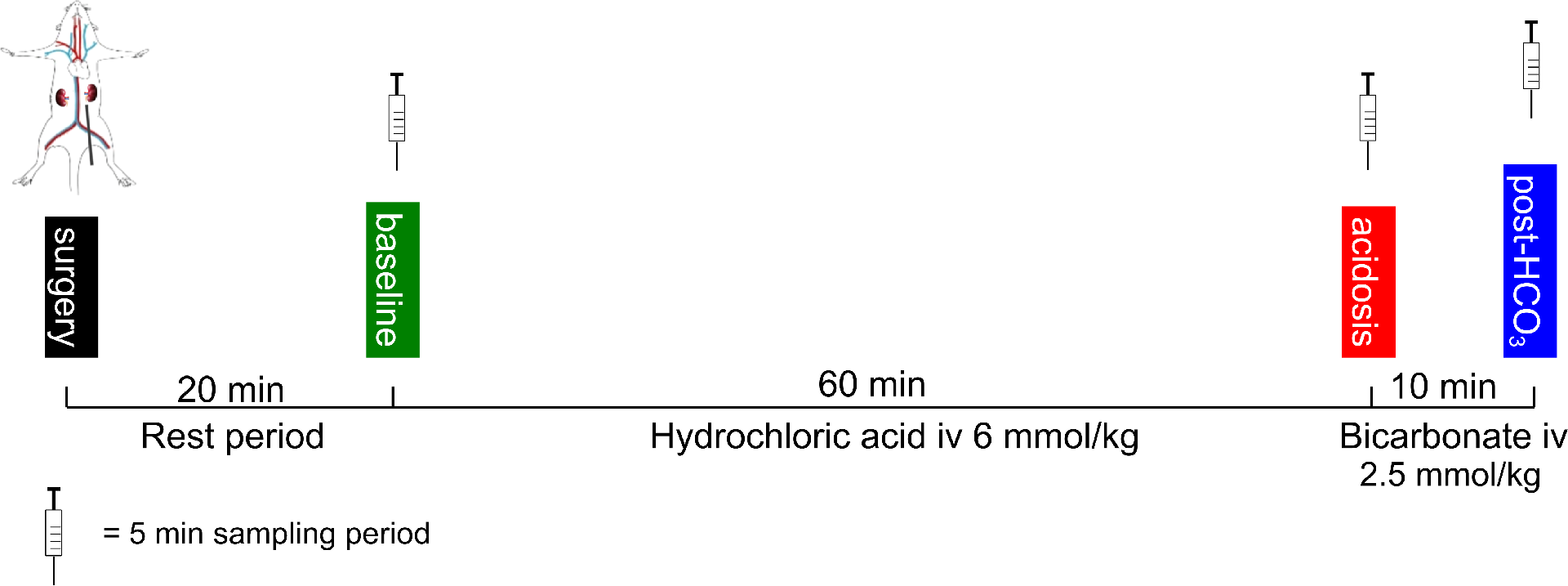
Experimental timeline. After an initial resting period of 20 minutes post-surgery, baseline measurements were made after which hydrochloric acid 6 mmol/kg was given as an infusion during a period of 60 min. New measurements were then performed after which sodium bicarbonate 2.5 mmol/kg was given as an intravenous infusion during a period of 10 minutes after which final measurements were performed.

### Glomerular transport studies

During the entire course of the experimental procedure, a continuous iv infusion (10 ml kg^−1^ h^−1^) of FITC-Ficoll (FITC-Ficoll-70, 20 μg ml-1; FITC-Ficoll-400, 480 μg ml-1; FITC-Inulin, 500 μg ml-1 and ^51^Cr-EDTA, 0.3 MBq ml^−1^) was given after an initial bolus dose (FITC-Ficoll-70, 40 μg; FITC-Ficoll-400, 960 μg; FITC-Inulin 0.5 mg and ^51^Cr-EDTA 0.3 MBq). This 1:24 mixture of Ficoll provides a broad range of molecular sizes. Sieving measurements were performed by a 5 min collection of urine from the left ureter with a mid-point (2.5 min) plasma sample. Glomerular filtration rate was assessed using the clearance of ^51^Cr-EDTA. The distributed pore model by Deen, Bridges, Brenner and Myers (11), modified by using two pore size distributions (31), was used to analyze the θ data for FITC-Ficoll (mol. radius 1.5–8.0 nm) by means of weighted nonlinear least squares regression to obtain the best curve fit as described previously (30). The solution of the model differential equations and calculation of renal plasma flow are described in (30).

### Determination of glomerular filtration rate in the left kidney

^51^Cr-EDTA (Amersham Biosciences, Buckinghamshire, UK) was given throughout the experiment *vide supra*. Radioactivity in blood (CPM_blood_) and urine (CPM_urine_) was detected using a gamma counter (Wizard 1480, LKP, Wallac, Turku, Finland). The glomerular filtration rate for the left kidney was estimated from the plasma to urine clearance of ^51^Cr-EDTA.

### Blood HCl titration experiments

Several arterial blood samples were obtained from one Sprague-Dawley rat and immediately analyzed with the I-STAT CHEM8+ cassette (Abbot Point of Care Inc, Abbot Park, IL) without and with the addition of 12 μmol HCl (Saltsyra APL 1 mmol/mL, APL, Stockholm, Sweden) per mL blood.

### High-performance size-exclusion chromatography (HPSEC)

To determine the concentration and size distribution of the FITC-Ficoll samples, a HPLC system (Waters, Milford, MA) was used. The plasma and urine samples were size-separated using an Ultrahydrogel 500 column (Waters) connected to a guard column (Waters) using a phosphate buffer (0.15 M NaCl, pH 7.4) as the mobile phase. The mobile phase was driven by a pump (Waters 1525). Fluorescence was detected at an excitation wavelength 492 nm and emission wavelength of 518 nm. An autosampler (Waters 717 plus) was used to load samples onto the system. The system was controlled by Breeze Software 3.3 (Waters). The system was calibrated using narrow Ficoll, dextran and protein standards as is described at some length in a previous article (3). The urine FITC-Ficoll concentration for each size (a_e_) was divided by the urine FITC-Inulin concentration to obtain the concentration of Ficoll in primary urine (C_u_). The glomerular sieving coefficient was then calculated by dividing C_u_ by the plasma concentration of Ficoll.

### Physiological acid-base parameters

The normal values assumed in the current work are shown in Supplemental Table 1. The concentration of titratable hydrogen ions (also called *base deficit*) in whole blood (*c*tH^+^_B_) and plasma (*c*tH^+^_P_) was calculated using the Van Slyke equation (see Supplemental material). The standard base excess (SBE) is the base excess of the extracellular fluid (6) and was calculated as follows

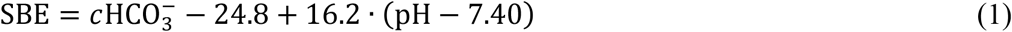

The concentration of titratable hydrogen ions in whole blood (*c*tH^+^_B_), also referred to as *blood base deficit*, was calculated using the following version (36) of the Van Slyke equation

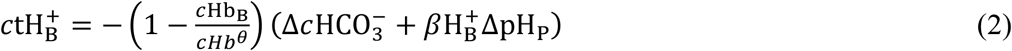

where *c*Hb_B_ (mmol/L) is the molar hemoglobin monomer concentration in blood (*c*Hb_B_ = *ρ*Hb_B_/16.114 where *ρ*Hb_B_ is the mass concentration of Hb in blood in g/L); *c*Hb^θ^ = 43 mmol/L is an empirical parameter accounting for the unequal distribution of hydrogen ions between erythrocytes and plasma; Δ*c*HCO_3_^−^ = *c*HCO_3_^−^ − 24.5; βH^+^_B_ = 2.3 × *c*Hb_B_ + β_P_ where 2.3 is the apparent buffer capacity of hemoglobin monomer *in vivo* and β_P_ = 7.7 mmol/L is the net buffer value of non-bicarbonate buffers for a normal plasma protein concentration; and, finally, ΔpH_P_ = pH_P_ – 7.4 is the difference between the actual and normal pH. We define the *actual* blood base excess (BE) as -*c*tH^+^_B_ and the actual plasma base excess (-*c*tH^+^_P_) as -*c*tH^+^_B_ when *c*Hb_B_=0. The normal values assumed in the current work are shown in Supplemental Table 1.

### Physiological model of erythrocyte and blood plasma acid-base titration

The blood plasma and red cell cytoplasm was modeled as an aqueous solution of polyprotic and monoprotic (Brønsted-Lowry) acids (H_n_A, H_m_B,…) which are partly dissociated into their conjugate bases as follows:

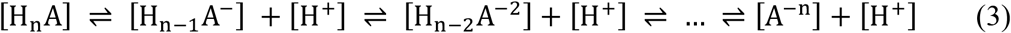

The total analytical concentration of an acid C_A_ can thus be written:

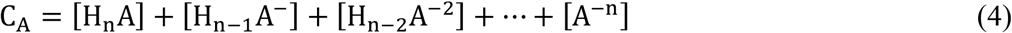

Each conjugate anion is associated with a corresponding dissociation coefficient:

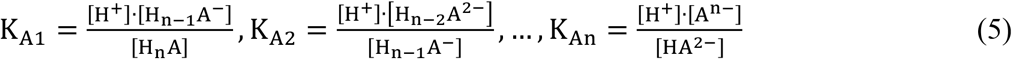

Using the dissociation coefficients we can re-write the acid concentration in terms of the hydrogen ion concentration and [H_n-1_A^−^] as follows:

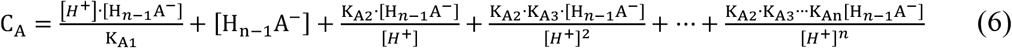

which is equivalent to:

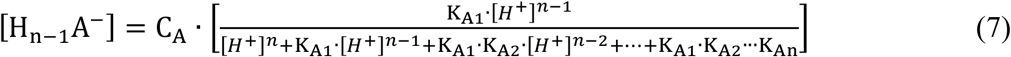

The total concentration C_A_ can thus be re-written in terms of [H_n-2_A^2−^], [H_n-3_A^3−^] into the well known (46) expression:

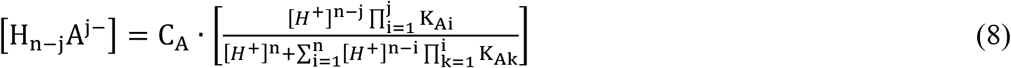

In equation S7, the product (Π) and the sum (Σ) operators have the following meanings: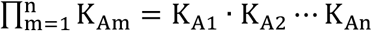 and 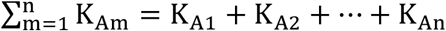 respectively.

Equation 7 is valid for all conjugate bases in the solution, which will equilibrate to the hydrogen ion concentration in an *isohydric* fashion. Thus, knowing the concentration of one acid and at least one of its conjugate bases will determine the hydrogen ion concentration (e.g. the Henderson-Hasselbalch equation). For carbonic acid (CA) the total concentration is as follows:

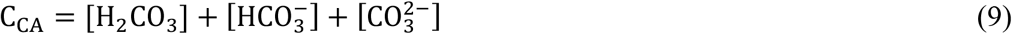

We used the following dissociation coefficients for CA:

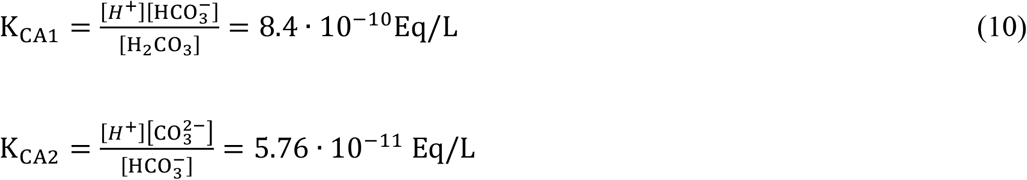

Using equation S7, we can write the first conjugate base of carbonic acid (i.e. bicarbonate) as follows

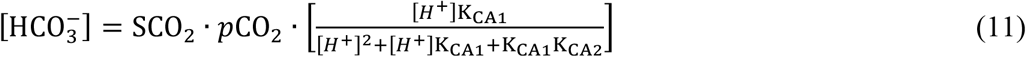

Similarly, the second conjugate base, i.e. carbonate, can be written as

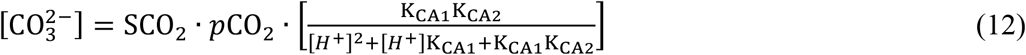

Here SCO_2_ is Henry’s law solubility coefficient for CO_2_, being ~0.225 mol/L/kPa (~0.03 mol/L/mmHg) at body temperature. Similarly, for phosphoric acid (PA):

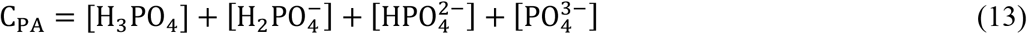

with dissociation coefficients:

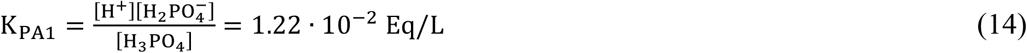

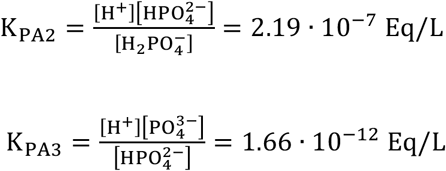

The conjugate bases can be written as follows:

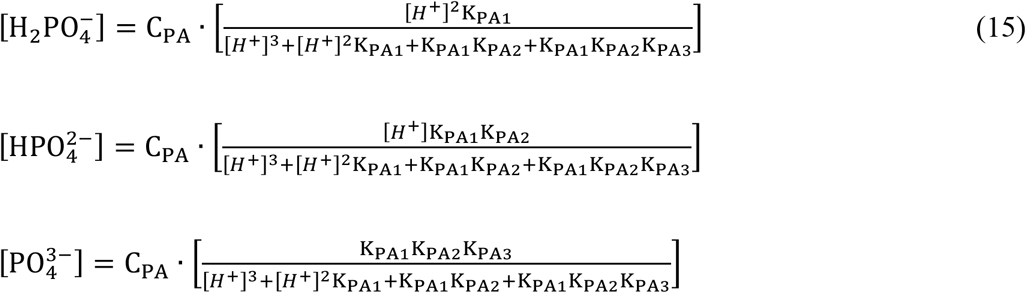

For serum albumin we used the model by Watson (44):

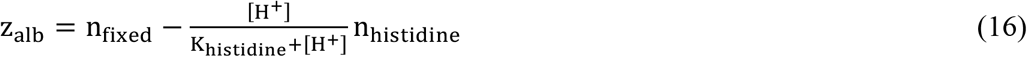

where n_histidine_ = 16 is the number of histidine residues on the albumin protein and n_fixed_ = 21 is the number of “fixed” negative charges on albumin during physiological conditions. For the histidine dissociation coefficient, we used K_histidine_ = 1.77·10−7 Eq/L. According to the *electroneutrality principle*, the total amount of charge in the red cell cytoplasm and in blood plasma (the ion mass balance) is zero:

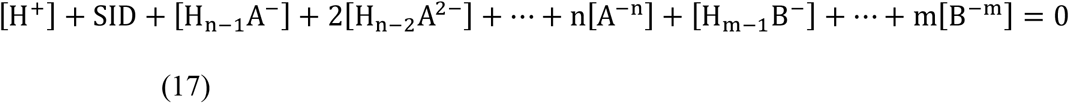

where SID is the strong ion difference, i.e. the sum of all *aprote* cations and anions in plasma:

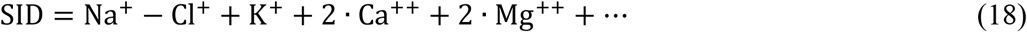

and so on. In our model of blood plasma we used:

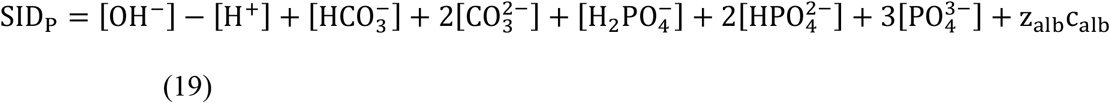

and in the erythrocyte:

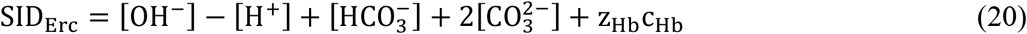

where we used the Hb titration equation applied in the model by Bookchin & Lew (26):

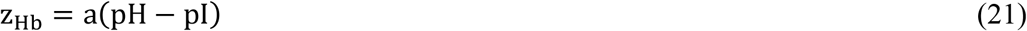

where pI=6.85 is the iso-electric point of hemoglobin and a = −10 mEq/pH unit. The erythrocyte hemoglobin tetramer concentration c_Hb_ was calculated using the molecular weight of hemoglobin, c_Hb_ = Hb/Hct/64,458 Da. For simulation purposes the albumin concentration was set to 42 g/L and the phosphate concentration to 1 mmol/L. Based on mass balance, the apparent base excess for the ECF taking the buffer effect of the erythrocytes into account can be calculated as follows:

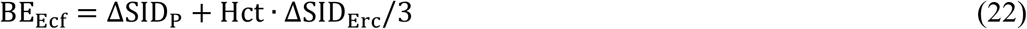

Here ΔSID_Erc_ and ΔSID_P_ is the difference in SID from a pH of 7.40 and a *p*CO2 of 5.33 kPa (40 mmHg). Similar to the SBE, this equation is based on the estimated hemoglobin concentration in the extracellular fluid (~1/3 of the blood hemoglobin concentration, see also ^2^). Model calculations were performed using R version 3.5.1 for macOS (The R Foundation for Statistical Computing).

### Statistical methods

The data are presented as median ± median absolute deviation unless otherwise specified. We used an alpha level (probability of type I error) of 0.05 and a beta level (probability of type II error) of 0.20 unless otherwise specified. A previous power analysis showed that four was the minimal number of animals needed to detect a difference in the sieving coefficient of Ficoll_70Å_ by a factor of at least 2 (12). Significant differences were assessed using an ANOVA on aligned rank transformed data (ARTool version 0.10.5), essentially performing a non-parametric test using parametric methods (45). We applied the modification of the F-statistic by Kenward and Roger (20) for small sample sizes. Tukey's Honest Significant Difference *post-hoc* tests were applied. Linear regression was performed using the *lm* function in R. Statistical analysis was performed using R version 3.5.1 for macOS (The R Foundation for Statistical Computing).

## RESULTS

### Severe acidemia induced by HCl and its correction using sodium bicarbonate does not alter glomerular permeability to Ficoll

Infusion of hydrochloric acid over the course of 60 minutes, 6 mmol/kg, evoked severe acidemia in Sprague-Dawley rats (n = 8), being partly improved by a 10 min iv infusion of sodium bicarbonate (0.6 mmol/mL) 2.5 mmol/kg (Figure 1 and Figure 2). The standard base excess (6), defined as the base excess (i.e. negative value denotes base deficit) of the extracellular fluid, decreased from 3 ± 1 mmol/L to −22 ± 2 mmol/L (Figure 2b). Assuming the extra-cellular volume (ECV) to be ~30% of the body weight, this corresponds to a H^+^ dose of ~20 mmol per L ECV, in concordance with the obtained result. Bicarbonate infusion (4.2 mL kg^−1^) apparently resulted in hemodilution (Figure 2c), corresponding to a ~15% increase of the plasma volume, whereas there were no significant alterations in hematocrit or hemoglobin level in acidemia compared to baseline (Figure 2c). Mean arterial pressure decreased from 125 ± 6 mmHg to 115 ± 9 mmHg (P = 0.02) in acidemia (Figure 2d). The partial CO_2_ pressure (*p*CO_2_) increased in acidemia, presumably due to buffer consumption (Figure 2e). Conceivably, since ventilator settings were set constant over the course of the experiment, a similar increment in *p*CO_2_ should occur after administration of bicarbonate, but the difference was non-significant (P = 0.06). The impact of acidemia on glomerular filtration rate was heterogeneous with some animals developing severe hypofiltration (Figure 2f). Plotted in Figure 3 are the glomerular sieving curves (θ *vs.* Stokes-Einstein radius) at baseline, before, and after bicarbonate infusion. Statistical comparison (Figure 3: bottom plot) of the sieving coefficients revealed no significant alterations in glomerular permeability, neither before nor after bicarbonate treatment. Using the distributed two-pore model, we noted no significant alterations except a small increment in the large pore radius post-HCO_3_^−^ (Table 2). Bicarbonate therapy lead to sodium loading (Figure 4a) (22) and a potassium shift into the intra-cellular compartment (Figure 4b) (1), decreasing K^+^ from 4.0 ± 0.3 mmol/L in acidosis to 3.5 ± 0.3 mmol/L after sodium bicarbonate, corresponding to a decrease in K^+^ of 0.4 mmol/L per 0.1 unit increase in pH. Following HCl infusion, plasma chloride increased from 101 ± 2 mmol/L to 121 ± 1 mmol/L, in good agreement with the administered dose of HCl (*vide infra*), and decreased to 116 ± 2 mmol/L after buffer infusion (Figure 4c). Blood urea nitrogen levels remained stable throughout the course of the experiment (Figure 4d).

**Figure 2.**
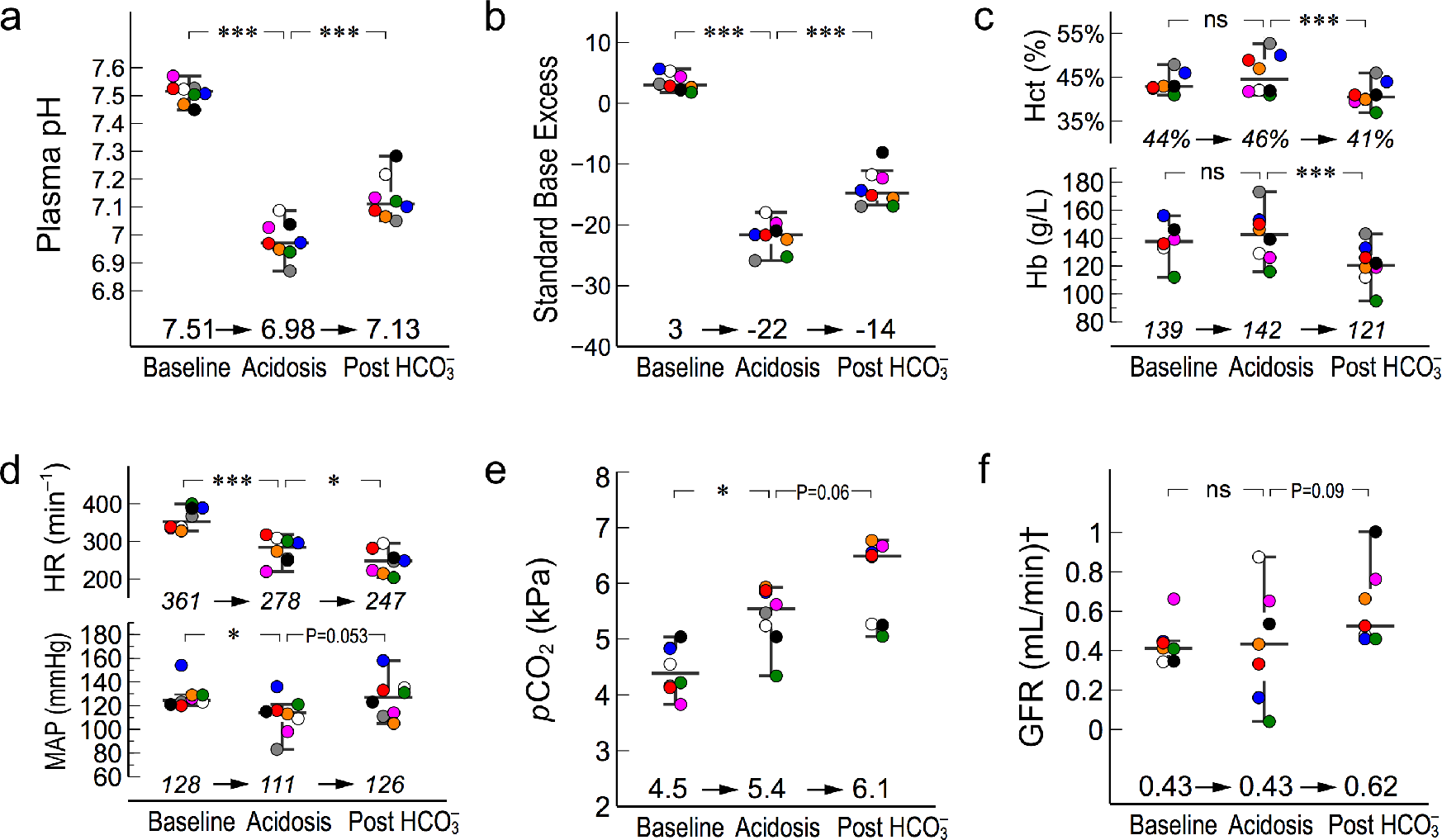
Infusion of hydrochloric acid (6 mmol/kg) evoked severe acidemia in Sprague-Dawley rats, being partly mitigated by sodium bicarbonate iv infusion (2.5 mmol/kg). Bicarbonate therapy improved the resulting base deficit by ~40% as quantified by the standard base excess (*a* and *b*). Buffer therapy resulted in hemodilution as indicated by a lower hematocrit (Hct) and hemoglobin (Hb) level (*c*). Mean arterial pressure and heart rate (HR) decreased significantly during acidemia, and HR decreased further after bicarbonate was given (*d*). The partial CO_2_ pressure increased in acidemia (*e*). The impact of acidemia on glomerular filtration was heterogeneous with some animals developing severe hypofiltration (*f*). † left kidney.

**Figure 3.**
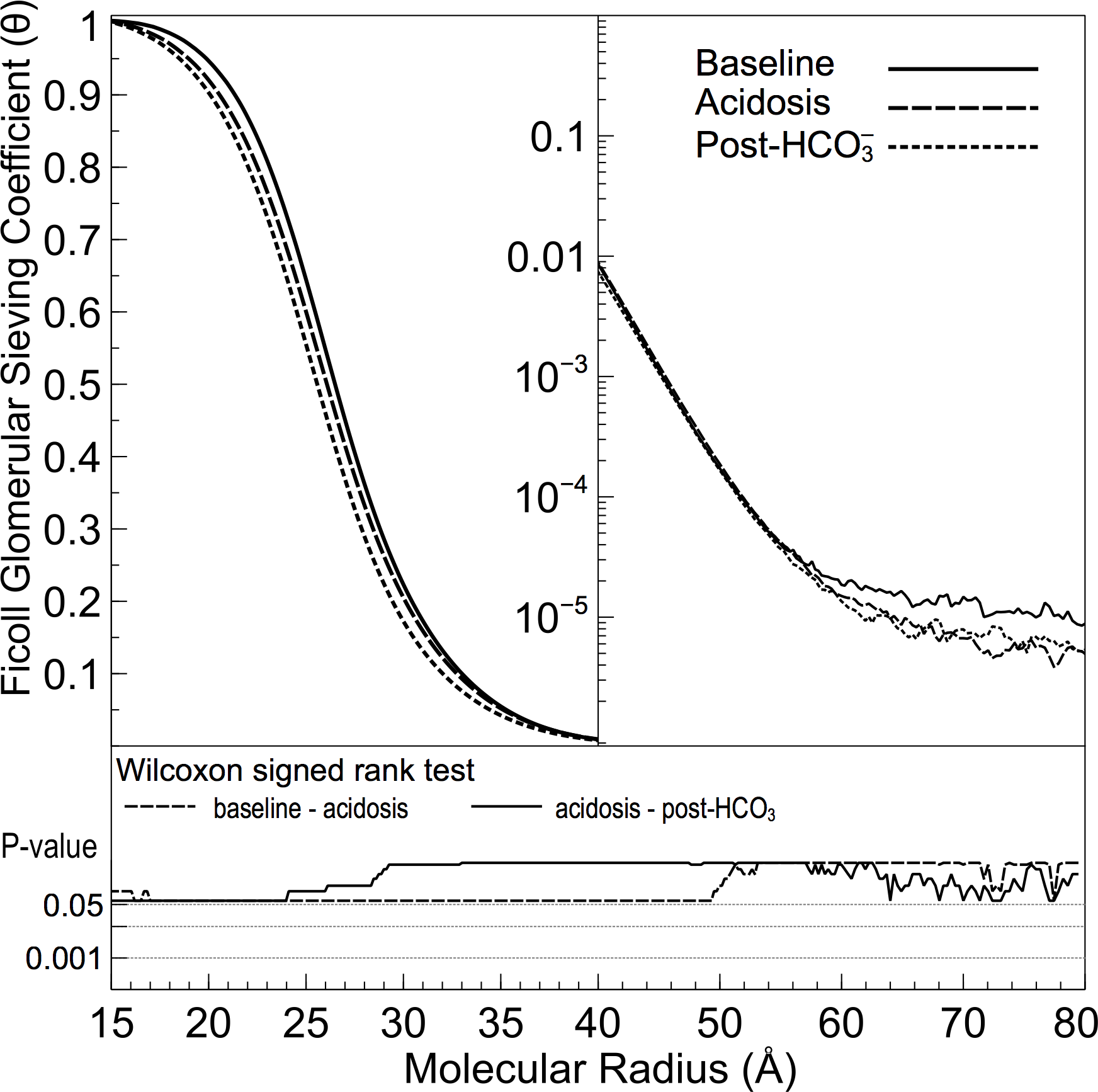
Glomerular sieving curves (Ficoll θ vs. hydrodynamic Stokes-Einstein radius) at baseline (solid line), before (dashed line) and after (dotted line) bicarbonate infusion. Statistical comparison (bottom plot) - across the range of the different molecular sizes – and between the different groups (baseline, before vs. after NaHCO3) revealed no significant alterations in glomerular permeability.

**Figure 4.**
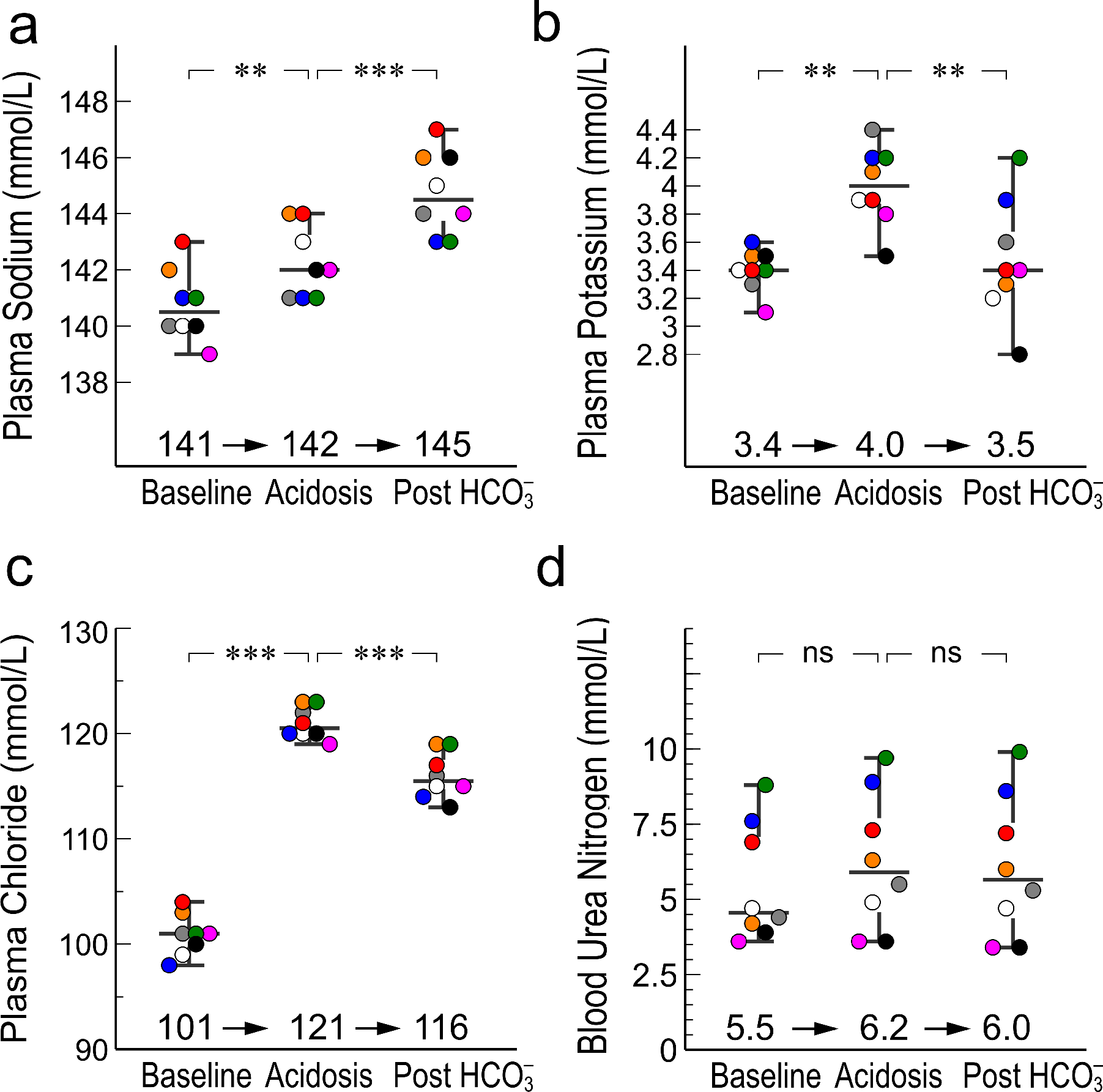
Electrolyte concentrations before and during acidemia, and after buffer therapy. Both HCl therapy and bicarbonate therapy lead to a significantly increased plasma sodium concentration (*a*), shifts in serum potassium from/to the intra-cellular compartment (*b*) (see also (1)), and large alterations in plasma chloride concentration (*c*). There were no significant changes in the plasma urea concentration during the experiment (*d*).

### Several established methods to assess buffer deficit are highly correlated but result in different estimates of the buffer deficit

Many different methods have been proposed to assess metabolic acid-base disorders and we find that the capability of several methods (Standard bicarbonate, total CO_2_ *et cetera*) to detect the existence of a significant acid-base disturbance is very similar (Supplemental Fig. 1). However, to guide buffer therapy it is also of clinical importance that a method can correctly identify the actual base deficit. In this regard, the various methods differed (Table 1) when compared to a calculated buffer deficit from the HCl dose assuming an ECV of 288 mL/kg (7) and no metabolic compensation.

**Table 1.** Predicted buffer deficit for different methods to assess acid-base status

| Method |  | Predicted buffer deficit |
| --- | --- | --- |
| BEFORE BUFFER THERAPY |  |  |
| *Calculated* from: | HCl dose (6 mmol/kg) † | 21 mmol/L |
| *Measured* using: | Standard Base Excess (SBE) | 22 ± 2 mmol/L<sup>a</sup> |
|  | Simplified ΔSID | 17 ± 2 mmol/L<sup>b</sup> |
|  | Standard Bicarbonate ‡ | 13 ± 2 mmol/L<sup>b</sup> |
|  | Total Carbon Dioxide * | 15 ± 1 mmol/L<sup>b</sup> |
| AFTER BUFFER THERAPY |  |  |
| *Calculated* from: | Bicarbonate dose (2.5 mmol/kg) † | 12 mmol/L |
| *Measured* using: | Standard Base Excess (SBE) | 14 ± 3 mmol/L<sup>c</sup> |
|  | Simplified ΔSID | 9 ± 1 mmol/L<sup>d</sup> |
|  | Standard Bicarbonate ‡ | 10 ± 2 mmol/L<sup>d</sup> |
|  | Total Carbon Dioxide * | 9 ± 1 mmol/L<sup>d</sup> |

† assuming an ECV of 288 mL/kg (cf. Bianchi et al (7)) and no metabolic compensation.
‡ assuming a normal value of 24.5 mmol/L.
\* assuming a normal value of 25.5 mmol/L.
<sup>a</sup> P=0.31 when compared to calculation using HCl dose.
<sup>b</sup> P<0.05 when compared to calculation using HCl dose.
<sup>c</sup> P=0.20 when compared to calculation using Bicarbonate dose.
<sup>d</sup> P<0.05 when compared to calculation using Bicarbonate dose.

**Table 2.** Two-pore model results

| Parameter | Baseline | Acidosis | Post-HCO<sub>3</sub><sup>-</sup> |  |
| --- | --- | --- | --- | --- |
| Small pore radius (Å) | 36 ± 1 | 35 ± 2 | 34 ± 1 | Small<br>Pore<br>Pathway |
| Small pore geometric SD | 1.15 ± 0.01 | 1.16 ± 0.02 | 1.16 ± 0.01 |  |
| A<sub>0</sub>/Δx × 10<sup>5</sup> (cm) † | 7.9 ± 0.4 | 8.2 ± 0.8 | 8.4 ± 0.4 |  |
| Large pore radius (Å) | 95 ± 16 | 76 ± 13 | 107 ± 9 ** | Large<br>Pore<br>Pathway |
| Large pore geometric SD | 1.03 ± 0.01 | 1.04 ± 0.02 | 1.05 ± 0.01 |  |
| α<sub>L</sub> × 10<sup>-5</sup> ‡ | 12 | 3 | 3 |  |
| Glomerular filtration pressure (mmHg) ¶ | 5 ± 1 | 6 ± 3 | 7 ± 2 | Hemodynamic<br>Parameters |
| GFR (mL/min) * | 0.4 ± 0.1 | 0.4 ± 0.2 | 0.6 ± 0.2 |  |
| Renal Plasma Flow (mL/min) <sup>*a</sup> | 1.3 ± 0.2 | 1.6 ± 0.7 | 2.0 ± 0.7 |  |
| Filtration Fraction (%) <sup>a</sup> | 32 ± 0 | 30 ± 3 | 23 ± 2 |  |

All values are median ± median absolute deviation.
† Unrestricted pore area (A<sub>0</sub>) over diffusion distance (Δx).
‡ Fractional large pore area (A<sub>0,L</sub>/A<sub>0</sub>)
¶ Calculated assuming a hydraulic conductance of 0.08 mL min
\* Left kidney only
\*\* P < 0.01 compared to acidosis phase.
<sup>a</sup> Calculated parameter (See Methods).

### The Simplified ΔSID Systematically Underestimates Buffer Deficit Assessed by SBE

Shown in Figure 5a is the simplified ΔSID = Na – Cl – 38, which decreased from 2 ± 1 at baseline to −17 ± 2 following HCl infusion, and increased to −9 ± 1 after bicarbonate therapy. Very similar alterations occurred for the plasma base excess (BE_P_; Figure 5b) and linear least-squares regression revealed a very high degree of correlation (R^2^ = 0.974), with nearly a 1:1 agreement, between ΔSID and BE_P_ (Figure 5c). Indeed, such a similitude is theoretically expected on the basis of the equality between true SID and plasma buffer base (36) and thus, it would appear from our data, that the simplified ΔSID is a strong predictor of the true change in SID. In contrast, inasmuch as the *standard* base excess (the amount of titrable acid in the extracellular fluid) and simplified ΔSID were highly correlated (R^2^ = 0.968), ΔSID *systematically underestimated* the buffer deficit assessed by standard base excess (SBE = 1.3543 × ΔSID + 0.1086, P < 0.001) (Figure 5d).

**Figure 5.**
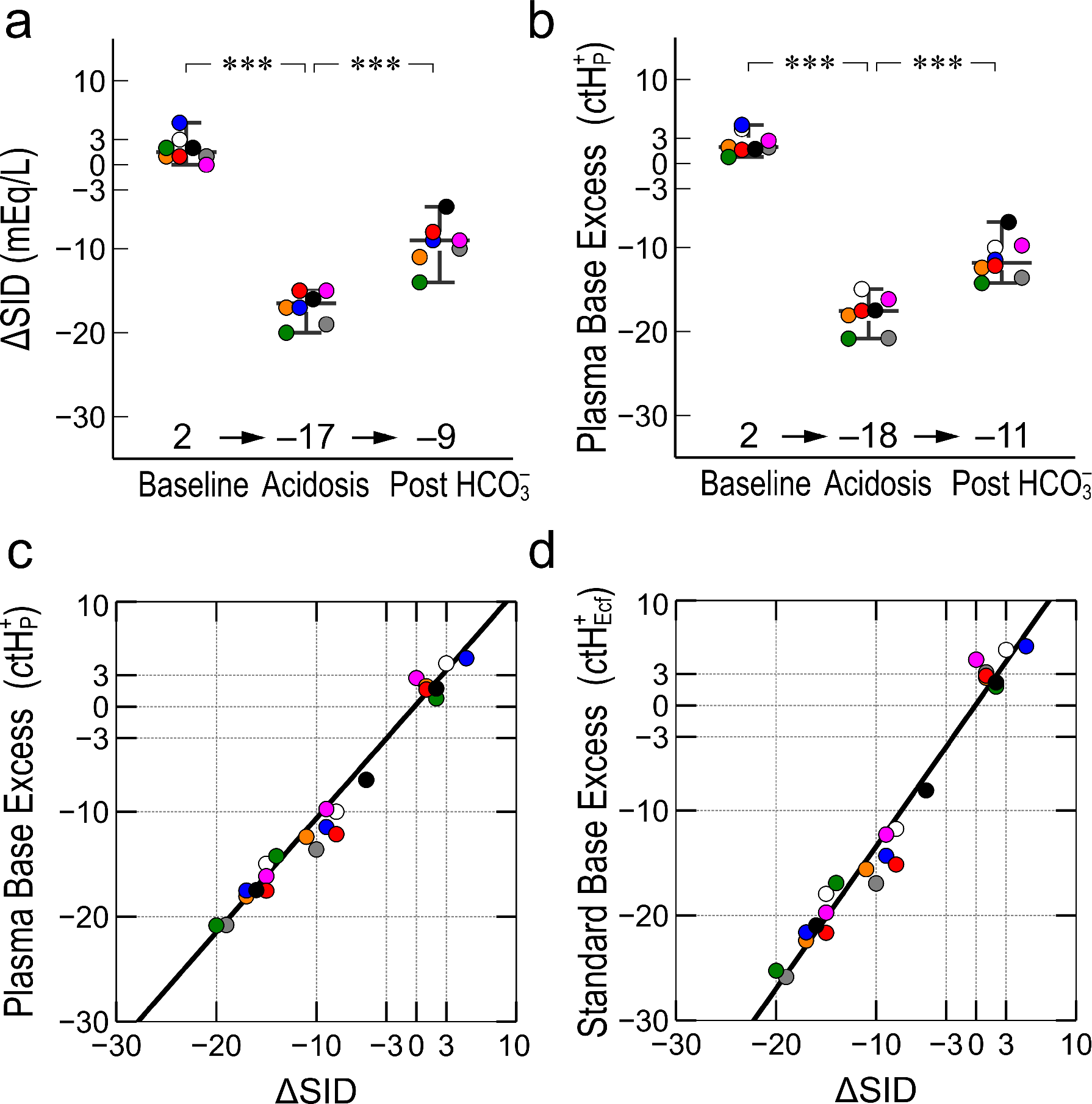
Infusion of hydrochloric acid (6 mmol/kg) decreased the simplified ΔSID = Na – Cl – 38, from 2 ± 1 at baseline to −17 ± 2, and increased to −9 ± 1 after bicarbonate therapy (a). Strikingly similar, and theoretically expected, alterations occurred for the plasma base excess BE_P_ (*b*) and linear regression unveiled an excellent agreement (R^2^ = 0.9742) between ΔSID and BE_P_ with a correlation coefficient close to unity (*c*). In contrast, although standard base excess and the simplified ΔSID were highly correlated (R^2^ = 0.9683), simplified ΔSID apparently underestimated the standard base excess (SBE = 1.3543 × ΔSID + 0.1086, P < 0.001) (*d*).

### Changes in the Strong Ion Difference in Erythrocytes Mediates a Buffering Effect on the Extracellular Fluid Which Explains the Difference Between Simplified ΔSID and SBE

It is well recognized that erythrocytes, mainly via their hemoglobin content, have a buffering effect on the extracellular fluid. The main difference between the above simplified ‘Fencl-Stewart’ ΔSID and the ‘Siggaard-Andersen’ SBE is that the former parameter does not take this buffer into account. Accordingly, in this study buffer deficit assessed with ΔSID was found to be consistently lower than SBE (Figure 5d). Thus, due to their buffer effect, erythrocytes would act to increase plasma strong ion difference (lowering ΔSID), presumably by lowering plasma chloride via the abundant and highly efficient AE1 (Band 3) HCO_3_^−^/Cl^−^ exchanger, present on the cell surface of erythrocytes (41). To quantitate this effect, we developed a theoretical model for the strong ion difference, or more precisely, the amount of titratable acid in the extracellular fluid and in erythrocytes based on the observation that the erythrocyte internal pH = 0.77 × external pH over a wide range of pH (41). The pH in erythrocytes is thus 7.25 at a normal blood plasma pH of 7.40 (41), which, if taking the most important intracellular buffers (bicarbonate and hemoglobin tetramer (41)) into account, corresponds to an erythrocyte SID of 38 mEq/L (per liter of cells). Our theoretical model is in a near 1:1 linear relationship with the simplified ΔSID (Figure 6a) if the effect of erythrocytes is excluded (i.e. taking only bicarbonate, albumin and phosphate buffers into account). In contrast, when effects of erythrocytes also are included, the model showed a near perfect agreement with SBE (Figure 6b). Thus, our *in silico* simulations predict that the effects of metabolic acidosis would strongly affect the SID of erythrocytes (Figure 7), giving rise to a buffering effect due to a shift of strong anions from plasma to erythrocytes, and increasing the SID of the extracellular fluid. This ‘anion shift’ should be much larger in vitro and, in a separate simple experiment, titration of whole blood with hydrochloric acid confirmed that a substantial amount of chloride is indeed shifted into the erythrocytes (Supplemental Figure 1).

**Figure 6.**
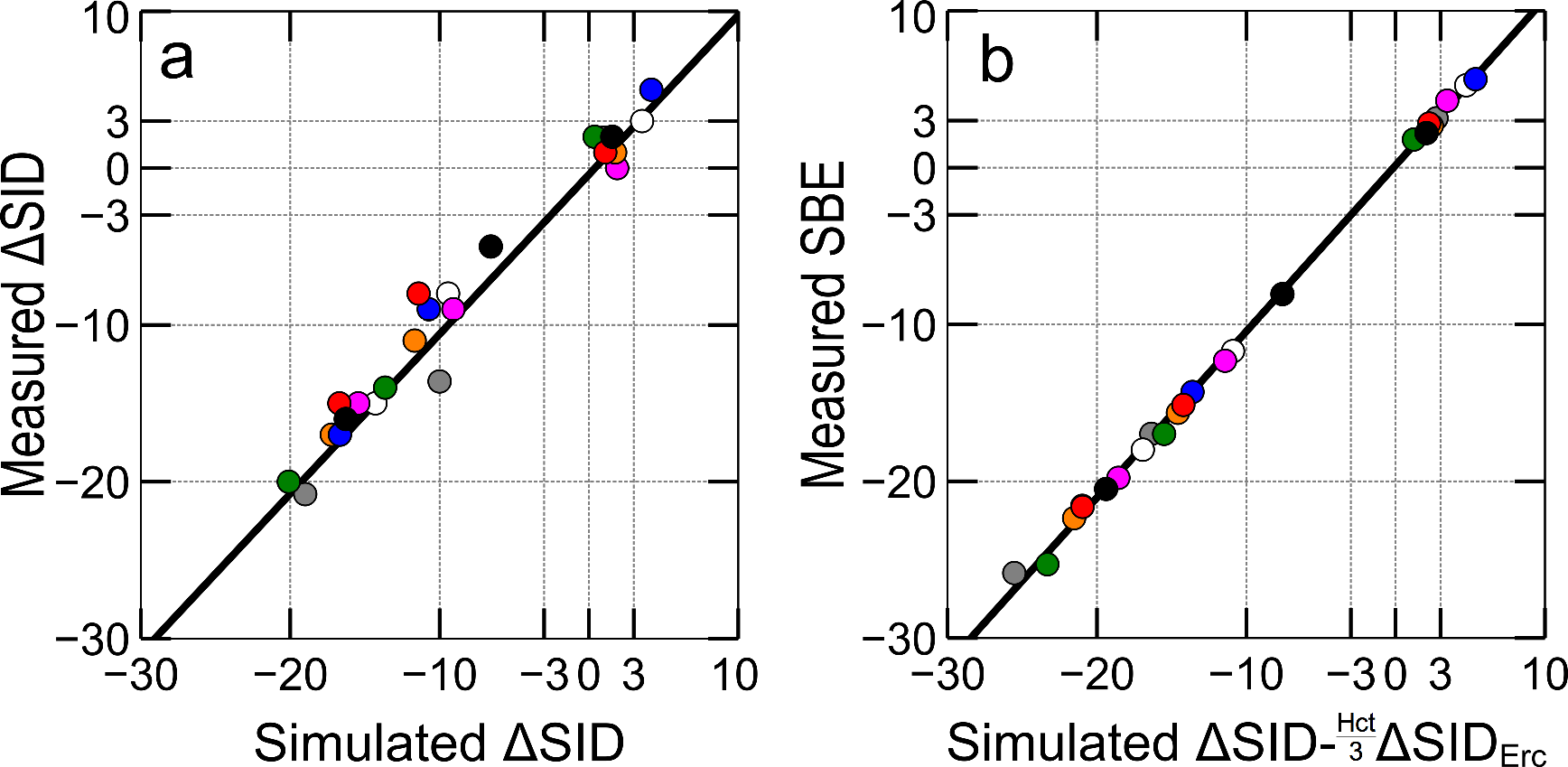
Linear regression between the theoretical model and experimental data: (*a*) Simulated extracellular fluid ΔSID *vs.* Measured simplified (Fencl-Stewart) ΔSID (R^2^ = 0.978, P < 0.001) and (*b*) Simulated extracellular fluid ΔSID corrected for the anion shift into erythrocytes compared with Measured Standard Base Excess (SBE) (R^2^ = 0.999, P < 0.001).

**Figure 7.**
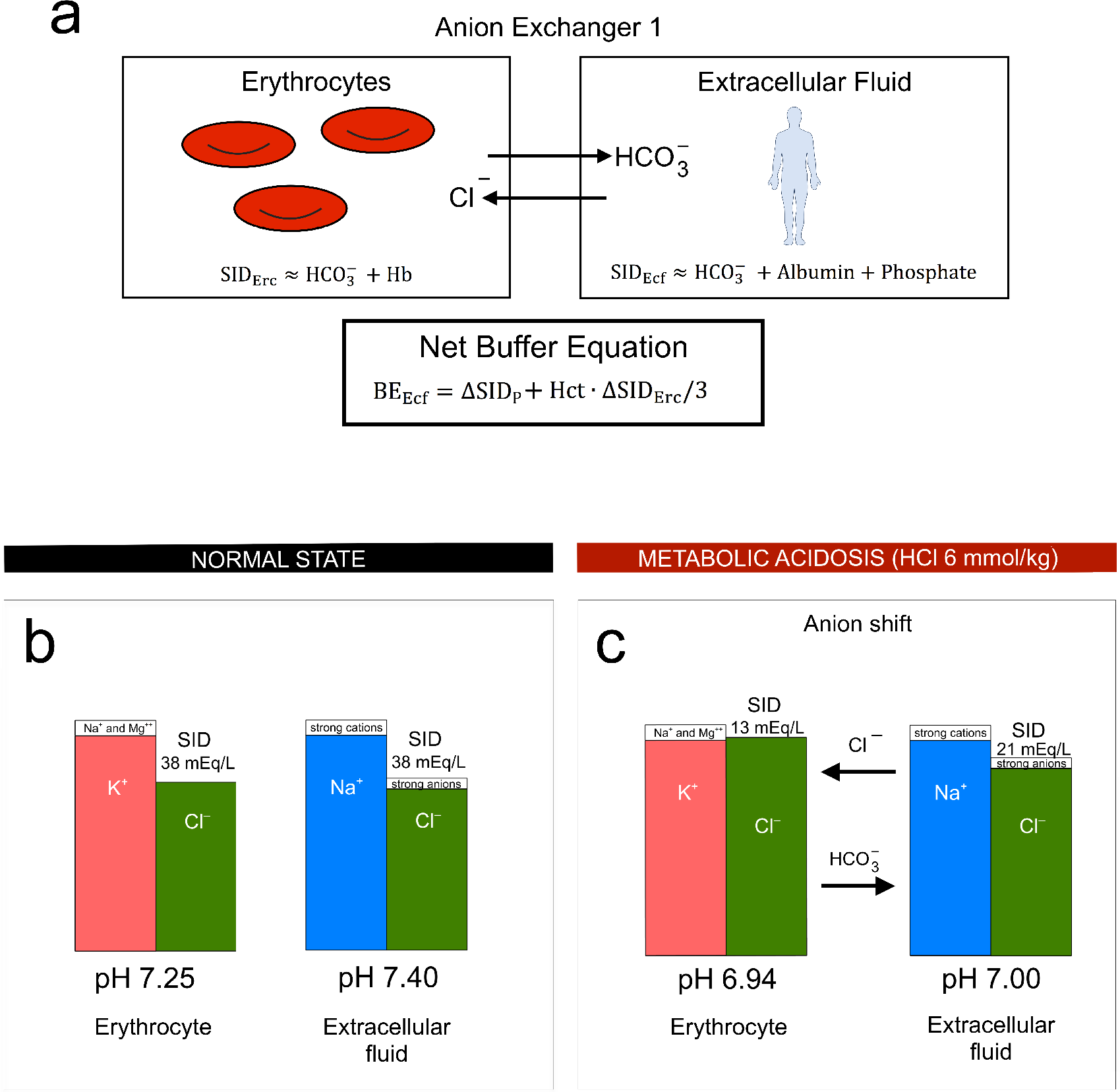
(*a*) Schematic representation of the theoretical model of the strong ion difference in the extracellular fluid and in erythrocytes: The buffering effect of erythrocytes on the extracellular fluid is mediated via a shift of anions from the extracellular fluid into the red cells, increasing the strong ion difference of the extracellular fluid. (*b*) The pH in erythrocytes is 7.25 at a normal blood plasma pH of 7.40 (41), which, if taking bicarbonate and hemoglobin tetramer into account, corresponds to an erythrocyte SID of 38 mEq per liter of cells (see Supplements). Erytrocyte buffer effect in severe acidosis (*b*): At a lower pH there will be a relatively higher increase in the anion concentration inside erythrocytes, leading to a lower anion concentration in extracellular fluid (thus increasing ΔSID).

## DISCUSSION

Emerging evidence unveils the crucial role of the cellular elements in the glomerular filtration barrier for the maintenance and regulation of glomerular permeability and function (12, 14, 32). Previous data from our group indicate that many disease mechanisms, including inflammatory stimuli (5, 40), angiotensin II (4), endothelin-1 (13), decreased NO bio-availability (14), ureteral obstruction (12) and hypoxia (34) lead to a glomerular hyperpermeability response. In light of these previous findings, it is surprising that no alterations in glomerular permeability were observed in the current study, with sieving coefficients for large Ficolls actually being numerically *lower* during severe acidemia.

An intracellular pH of 7.10-7.30 is essential for the normal function of cells (27) and it is commonly assumed that intracellular acidosis will occur whenever there is an extracellular acidosis and that it is the former that leads to the negative effects (24). There is however uncertainty and some controversy over whether the intra-cellular pH decrease occurs uniformly in all tissues and whether these alterations are sustained or transient and/or depend on the mechanism causing the acidosis (24). Indeed, other *in vivo* studies of acidosis due to HCl (or lactic acid) infusion have shown little or no effect on intra-cellular pH (48), while ischemic lactic acidosis caused marked effects on intra-cellular pH (33). Also, Shapiro *et al* (35) found that NaHCO_3_ treatment caused intracellular brain acidification in rats subjected to ammonium chloride acidosis. However, Bollaert et al (8) found no intra-cellular acidosis in the muscle tissue in septic rats with lactic acidosis after correction with NaHCO_3_ (35). Similarly, in an elegant study, Levraut and colleagues (25) demonstrated that intra-cellular acidosis only occurred *in vitro* when using a non-bicarbonate buffering system. Albeit, from a mechanistic point of view, increasing the pH of both compartments should be beneficial in comparison with just increasing the extracellular pH. Thus, it may at first seem obvious that sodium bicarbonate therapy would be beneficial in the treatment of acidosis. However, the potential benefit of buffer therapy has been a matter of much debate. For example, nephrologists and critical care physicians had a clear disparity of opinion when polled about the use of buffer therapy in severe organic acidosis (21). While the beneficial effects of sodium bicarbonate on hemodynamics are robustly supported by experimental studies, its ability to improve hemodynamics and morbidity in critically ill patients is as yet unproven. In a very recent report (17), Jaber and colleagues randomized patients with pH ≤ 7.20 to either NaHCO_3_ to maintain a pH ≥ 7.30 or no treatment and found no difference in the primary outcome of mortality at day 28 or organ failure at day 7. However, in a predefined stratum of patients with acute kidney injury network (AKIN) scores of 2-3, mortality was less frequent in the bicarbonate group compared to controls. Furthermore, the need for renal replacement therapy was overall lower in the bicarbonate group (17).

A common indication for buffer therapy is hemodynamic instability in severe acidemia. In addition, the condition of critically ill patients may be further complicated by acidemia-induced hyporesponsiveness to vasopressors. It is also well known that acidemia affects the potency and risk of side effects of numerous drugs used in anesthesia and intensive care. For example, volatile anesthetics become more arrhythmogenic while opiates and neuromuscular blockers are potentiated. In the current study we found a decreased mean arterial pressure and heart rate during acidosis, but our data failed to show improvements following bicarbonate therapy. However, in mild to moderate acidemia, pH > 7.20, cardiac output can actually increase due to enhanced catecholamine release (22), but even mild acidemia has been associated with hemodynamic instability (19, 22). For example, Kellum and colleagues induced different degrees of hyperchloremic acidosis in septic rats and demonstrated a significantly lower arterial pressure in the animals with mild acidemia (SBE −5 to −10) compared with non-acidotic animals (19). However, neither controlled clinical studies (10, 28) nor experimental studies (9) in lactic acidosis showed any benefit on cardiovascular function following NaHCO_3_ administration.

Acute and chronic metabolic acidosis are common acid–base disorders and their diagnosis relies on the correct estimation of the buffer deficit, identification of respiratory compensation, and an assessment of the plasma anion gap, preferably corrected according to the plasma albumin concentration (6) so as to be able to identify the etiology of the disorder. Here we provide evidence of significant differences between the standard base excess compared to other methods. A priori, this result can be expected since none of the other methods include the buffering effect of erythrocytes. Furthermore, the current results highlight the difference between the quantification of buffer deficit and the assessment of ion mass balance, e.g. in the form of the anion gap. Mathematically, the albumin corrected anion gap is identical to using the simplified ΔSID and adding 1 mEq/L per every 4 g decrease in plasma albumin from a normal concentration of 42 g/L (6, 39). The clinical rationale for such computations is to detect organic, anion-gap acidosis (6). While the use of only sodium and chloride and bicarbonate/base excess to assess ion mass balance may seem simplistic compared to summing up all plasma cations and anions, one should bear in mind that including more measurements will inflate the coefficient of variation of the composite parameter and, thus, decrease precision. Indeed, our current data show that the simplified ΔSID = Na – Cl – 38 can predict the actual ΔSID with surprising accuracy.

We developed a strong ion theory similar to that by Stewart (37), but also including the buffering effect of erythrocytes. We show that this modification leads to the near equality of the buffer deficit approximated by SBE and that approximated with the novel model (Supplemental Equation 21). Our modelization rely heavily on the assumption that most of the buffering effect is mediated via the AE1 isoform of the anion-exchange transporter (also called Band 3), being highly expressed on erythrocytes with more than 10^6^ copies per red cell (41). It cannot however be excluded that other transport proteins contribute. For example, the sodium concentration also increased in our *in vitro* experiment (see Supplemental Figure 2), which may imply a role for the sodium-hydrogen anti-porter (NHE-1) in erythrocytes. Many other simplifying assumptions were made in our model. For example, we assumed that the only non-carbonate buffers are albumin and phosphate in plasma; and bicarbonate and hemoglobin in erythrocytes. We also used a simplistic equation for hemoglobin titration. Indeed, far more sophisticated models have been developed for this purpose (47). Nevertheless, despite its simplicity, we find that our model provides an excellent fit to our data. It also serves to convey a fundamental understanding, and simple quantitation, of the buffering effects of erythrocytes.

We conclude that neither acidosis nor its correction using sodium bicarbonate appear to have any negative effects on glomerular permeability. Furthermore, the striking agreement between the plasma base excess and the simplified ΔSID supports the use of the simplified Fencl-Stewart approach (39). Inasmuch as the difference between SBE and ΔSID was significant, and most likely caused by the fact that SBE also takes erythrocyte buffering into account, the differences were rather small from a clinical perspective, even in a very severe acid-base disturbance such as in the present study. Thus, both methods should provide reasonable results in clinical practice. We conclude that our strong ion modelization may provide part of the “missing link” between strong ion theory and the Danish base excess concept. It however deserves mentioning that, in the present study, these similarities were shown in Sprague-Dawley rats and not in patients, and clinical studies could be performed to confirm or refute the present results. Regardless of with what method one prefers to assess acid-base status, it appears that the standard base excess, robustly supported by several clinical studies (6), is a more suitable method to determine the extent of a metabolic acid–base disorder while the simplified Fencl-Stewart approach (39), or the albumin corrected anion gap (6), aids the clinician in identifying the mechanism.

## AUTHOR CONTRIBUTIONS

J.S. and C.M.O. designed the study; A.R. and C.M.O. carried out experiments; C.M.O and J.S. analyzed the data; C.M.O. made the figures; C.M.O, P.B., D.P., N.C. and J.S. drafted and revised the paper; all authors approved the final version of the manuscript.

## ACKNOWLEDGEMENTS

This work was funded by the medical faculty in Lund (ALF grant). The authors declare no competing interests pertaining to the current article.

## REFERENCES

1. Adrogue HJ, and Madias NE. PCO2 and [K+]p in metabolic acidosis: certainty for the first and uncertainty for the other. J Am Soc Nephrol 15: 1667–1668, 2004.

2. Aronson PS, and Giebisch G. Effects of pH on potassium: new explanations for old observations. J Am Soc Nephrol 22: 1981–1989, 2011.

3. Asgeirsson D, Venturoli D, Rippe B, and Rippe C. Increased glomerular permeability to negatively charged Ficoll relative to neutral Ficoll in rats. Am J Physiol Renal Physiol 291: F1083–1089, 2006.

4. Axelsson J, Rippe A, Öberg CM, and Rippe B. Rapid, dynamic changes in glomerular permeability to macromolecules during systemic angiotensin II (ANG II) infusion in rats. Am J Physiol Renal Physiol 303: F790–799, 2012.

5. Axelsson J, Rippe A, Venturoli D, Swärd P, and Rippe B. Effects of early endotoxemia and dextran-induced anaphylaxis on the size selectivity of the glomerular filtration barrier in rats. Am J Physiol Renal Physiol 296: F242–248, 2009.

6. Berend K. Diagnostic Use of Base Excess in Acid-Base Disorders. N Engl J Med 378: 1419–1428, 2018.

7. Bianchi M, Bellini G, Hessan H, Kim KE, Swartz C, and Fernandes M. Body fluid volumes in the spontaneously hypertensive rat. Clin Sci (Lond) 61: 685–691, 1981.

8. Bollaert PE, Robin-Lherbier B, Mallie JP, Nace L, Escanye JM, and Larcan A. Effects of sodium bicarbonate on striated muscle metabolism and intracellular pH during endotoxic shock. Shock 1: 196–200, 1994.

9. Cooper DJ, Herbertson MJ, Werner HA, and Walley KR. Bicarbonate Does Not Increase Left-Ventricular Contractility during L-Lactic Acidemia in Pigs. Am Rev Respir Dis 148: 317–322, 1993.

10. Cooper DJ, Walley KR, Wiggs BR, and Russell JA. Bicarbonate does not improve hemodynamics in critically III patients who have lactic acidosis: A prospective, controlled clinical study. Annals of internal medicine 112: 492–498, 1990.

11. Deen WM, Bridges CR, Brenner BM, and Myers BD. Heteroporous model of glomerular size selectivity: application to normal and nephrotic humans. Am J Physiol 249: F374–389, 1985.

12. Dolinina J, Rippe A, Bentzer P, and Öberg CM. Glomerular hyperpermeability after acute unilateral ureteral obstruction: effects of Tempol, NOS, RhoA, and Rac-1 inhibition. Am J Physiol Renal Physiol 315: F445–F453, 2018.

13. Dolinina J, Rippe A, and Öberg CM. Sustained, delayed and small increments in glomerular permeability to macromolecules during systemic endothelin-1 infusion mediated via the ETA receptor. Am J Physiol Renal Physiol 2019.

14. Dolinina J, Sverrisson K, Rippe A, Öberg CM, and Rippe B. Nitric oxide synthase inhibition causes acute increases in glomerular permeability in vivo, dependent upon reactive oxygen species. Am J Physiol Renal Physiol 311: F984–F990, 2016.

15. Gardner KD, Jr. The effect of pH on the filtration, reabsorption, and excretion of protein by the rat kidney. J Clin Invest 40: 525–535, 1961.

16. Ilatovskaya DV, Palygin O, Levchenko V, Endres BT, and Staruschenko A. The Role of Angiotensin II in Glomerular Volume Dynamics and Podocyte Calcium Handling. Sci Rep 7: 299, 2017.

17. Jaber S, Paugam C, Futier E, Lefrant JY, Lasocki S, Lescot T, Pottecher J, Demoule A, Ferrandiere M, Asehnoune K, Dellamonica J, Velly L, Abback PS, de Jong A, Brunot V, Belafia F, Roquilly A, Chanques G, Muller L, Constantin JM, Bertet H, Klouche K, Molinari N, Jung B, and Group B-IS. Sodium bicarbonate therapy for patients with severe metabolic acidaemia in the intensive care unit (BICAR-ICU): a multicentre, open-label, randomised controlled, phase 3 trial. Lancet 392: 31–40, 2018.

18. Jung B, Rimmele T, Le Goff C, Chanques G, Corne P, Jonquet O, Muller L, Lefrant JY, Guervilly C, Papazian L, Allaouchiche B, Jaber S, and AzuRea G. Severe metabolic or mixed acidemia on intensive care unit admission: incidence, prognosis and administration of buffer therapy. A prospective, multiple-center study. Crit Care 15: R238, 2011.

19. Kellum JA, Song M, and Venkataraman R. Effects of hyperchloremic acidosis on arterial pressure and circulating inflammatory molecules in experimental sepsis. Chest 125: 243–248, 2004.

20. Kenward MG, and Roger JH. Small sample inference for fixed effects from restricted maximum likelihood. Biometrics 53: 983–997, 1997.

21. Kraut JA, and Kurtz I. Use of base in the treatment of acute severe organic acidosis by nephrologists and critical care physicians: results of an online survey. Clin Exp Nephrol 10: 111–117, 2006.

22. Kraut JA, and Madias NE. Metabolic acidosis: pathophysiology, diagnosis and management. Nat Rev Nephrol 6: 274–285, 2010.

23. Kraut JA, and Madias NE. Sodium bicarbonate for severe metabolic acidaemia. Lancet 392: 3–4, 2018.

24. Kraut JA, and Madias NE. Treatment of acute metabolic acidosis: a pathophysiologic approach. Nat Rev Nephrol 8: 589–601, 2012.

25. Levraut J, Labib Y, Chave S, Payan P, Raucoules-Aime M, and Grimaud D. Effect of sodium bicarbonate on intracellular pH under different buffering conditions. Kidney Int 49: 1262–1267, 1996.

26. Lew VL, and Bookchin RM. Volume, pH, and ion-content regulation in human red cells: analysis of transient behavior with an integrated model. J Membr Biol 92: 57–74, 1986.

27. Madshus IH. Regulation of intracellular pH in eukaryotic cells. Biochem J 250: 1–8, 1988.

28. Mathieu D, Neviere R, Billard V, Fleyfel M, and Wattel F. Effects of bicarbonate therapy on hemodynamics and tissue oxygenation in patients with lactic acidosis: a prospective, controlled clinical study. Crit Care Med 19: 1352–1356, 1991.

29. Mitchell JH, Wildenthal K, and Johnson RL, Jr. The effects of acid-base disturbances on cardiovascular and pulmonary function. Kidney Int 1: 375–389, 1972.

30. Öberg CM, Groszek JJ, Roy S, Fissell WH, and Rippe B. A distributed solute model: an extended two-pore model with application to the glomerular sieving of Ficoll. Am J Physiol Renal Physiol ajprenal000662017, 2017.

31. Öberg CM, and Rippe B. A distributed two-pore model: theoretical implications and practical application to the glomerular sieving of Ficoll. American journal of physiology Renal physiology 306: F844–854, 2014.

32. Perico L, Conti S, Benigni A, and Remuzzi G. Podocyte-actin dynamics in health and disease. Nat Rev Nephrol 12: 692–710, 2016.

33. Rehring TF, Shapiro JI, Cain BS, Meldrum DR, Cleveland JC, Harken AH, and Banerjee A. Mechanisms of pH preservation during global ischemia in preconditioned rat heart: roles for PKC and NHE. Am J Physiol 275: H805–813, 1998.

34. Rippe C, Rippe A, Larsson A, Asgeirsson D, and Rippe B. Nature of glomerular capillary permeability changes following acute renal ischemia-reperfusion injury in rats. Am J Physiol Renal Physiol 291: F1362–1368, 2006.

25. Shapiro JI, Whalen M, Kucera R, Kindig N, Filley G, and Chan L. Brain pH responses to sodium bicarbonate and Carbicarb during systemic acidosis. The American journal of physiology 256: H1316–1321, 1989.

36. Siggaard-Andersen O, and Fogh-Andersen N. Base excess or buffer base (strong ion difference) as measure of a non-respiratory acid-base disturbance. Acta Anaesthesiol Scand Suppl 107: 123–128, 1995.

37. Stewart PA. Modern quantitative acid-base chemistry. Can J Physiol Pharmacol 61: 1444–1461, 1983.

38. Story DA. Bench-to-bedside review: a brief history of clinical acid-base. Crit Care 8: 253–258, 2004.

39. Story DA, Morimatsu H, and Bellomo R. Strong ions, weak acids and base excess: a simplified Fencl-Stewart approach to clinical acid-base disorders. Br J Anaesth 92: 54–60, 2004.

40. Sverrisson K, Axelsson J, Rippe A, Asgeirsson D, and Rippe B. Acute reactive oxygen species (ROS)-dependent effects of IL-1beta, TNF-alpha, and IL-6 on the glomerular filtration barrier (GFB) in vivo. Am J Physiol Renal Physiol 309: F800–806, 2015.

41. Swietach P, Tiffert T, Mauritz JM, Seear R, Esposito A, Kaminski CF, Lew VL, and Vaughan-Jones RD. Hydrogen ion dynamics in human red blood cells. J Physiol 588: 4995–5014, 2010.

42. Thode J, Holmegaard SN, Transbøl I, Fogh-Andersen N, and Siggaard-Andersen O. Adjusted ionized calcium (at pH 7.4) and actual ionized calcium (at actual pH) in capillary blood compared for clinical evaluation of patients with disorders of calcium metabolism. Clin Chem 36: 541–544, 1990.

43. Throssell D, Harris KP, Bevington A, Furness PN, Howie AJ, and Walls J. Renal effects of metabolic acidosis in the normal rat. Nephron 73: 450–455, 1996.

44. Watson PD. Modeling the effects of proteins on pH in plasma. J Appl Physiol (1985) 86: 1421–1427, 1999.

45. Wobbrock JO, Findlater L, Gergle D, and Higgins JJ. The aligned rank transform for nonparametric factorial analyses using only anova procedures. In: Proceedings of the SIGCHI conference on human factors in computing systemsACM, 2011, p. 143–146.

46. Wooten EW. Analytic calculation of physiological acid-base parameters in plasma. J Appl Physiol (1985) 86: 326–334, 1999.

47. Wooten EW. Calculation of physiological acid-base parameters in multicompartment systems with application to human blood. J Appl Physiol (1985) 95: 2333–2344, 2003.

48. Zahler R, Barrett E, Majumdar S, Greene R, and Gore JC. Lactic acidosis: effect of treatment on intracellular pH and energetics in living rat heart. Am J Physiol 262: H1572–1578, 1992.

