## Supplemental Material for "Buffer Therapy in Acute Metabolic Acidosis: Effects on Glomerular Permeability and Acid-Base Status"

*[1] Department of Nephrology, Skåne University Hospital, Clinical Sciences Lund, Lund University, Lund, Sweden. [2] Department of Anesthesiology and Intensive Care, Clinical Sciences Lund, Lund University, Lund, Sweden. [3] Department of Anesthesia and Intensive Care, Helsingborg Hospital, Helsingborg, Sweden. [4] Department of Anesthesia and Intensive Care, Skåne University Hospital, Lund/Malmö, Sweden*

### SUPPLEMENTAL MATERIAL

Supplemental Figure 1 ………………………………………. page 2

Supplemental Figure 2 ………………………………………. page 3

Supplemental Table 1 ………………………………………. page 4

References ……………………………………………………….. page 5

**Supplemental Figure 1**

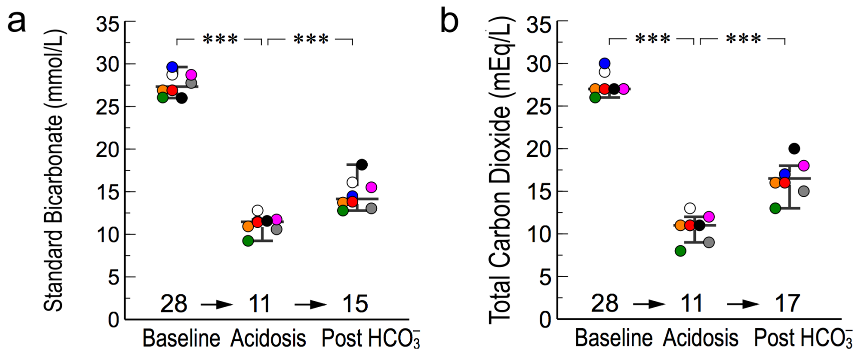


c

| | dSID | BE | SBE | TCO2 | HCO3 | StdBic | Cl |
|---|---|---|---|---|---|---|---|
| dSID | 1 | 0.98 | 0.98 | 0.99 | 0.99 | 0.97 | -0.97 |
| BE | 0.98 | 1 | 1 | 0.99 | 1 | 0.99 | -0.98 |
| SBE | 0.98 | 1 | 1 | 1 | 1 | 0.99 | -0.98 |
| TCO2 | 0.99 | 0.99 | 1 | 1 | 1 | 0.99 | -0.98 |
| HCO3 | 0.99 | 1 | 1 | 1 | 1 | 0.99 | -0.98 |
| StdBic | 0.97 | 0.99 | 0.99 | 0.99 | 0.99 | 1 | -0.99 |
| Cl | -0.97 | -0.98 | -0.98 | -0.98 | -0.98 | -0.99 | 1 |

Colour scale: 1, 0.8, 0.6, 0.4, 0.2, 0, -0.2, -0.4, -0.6, -0.8, -1

***Older methods to assess acid-base disorders*** include the *standard bicarbonate* (StdBic) introduced by Jørgensen and Astrup [1], defined as the plasma bicarbonate concentration at a *p*CO2 of 5.3 kPa (40 mmHg), and the *total plasma carbon dioxide concentration* (TCO2) defined as the actual bicarbonate concentration + 0.225 × $pCO_2$. Both these parameters were altered in similar ways before and after buffer loading (*a* and *b*). Subsequent correlation analysis showed that blood base excess (BE), standard base excess (SBE), simplified ΔSID (dSID), plasma chloride concentration (Cl) and the actual plasma bicarbonate concentration are highly correlated (*c*). Even just using the plasma chloride concentration (*c*) correlated well with SBE, which is expected in a non-anion gap acidosis.

**Supplemental Figure 2**

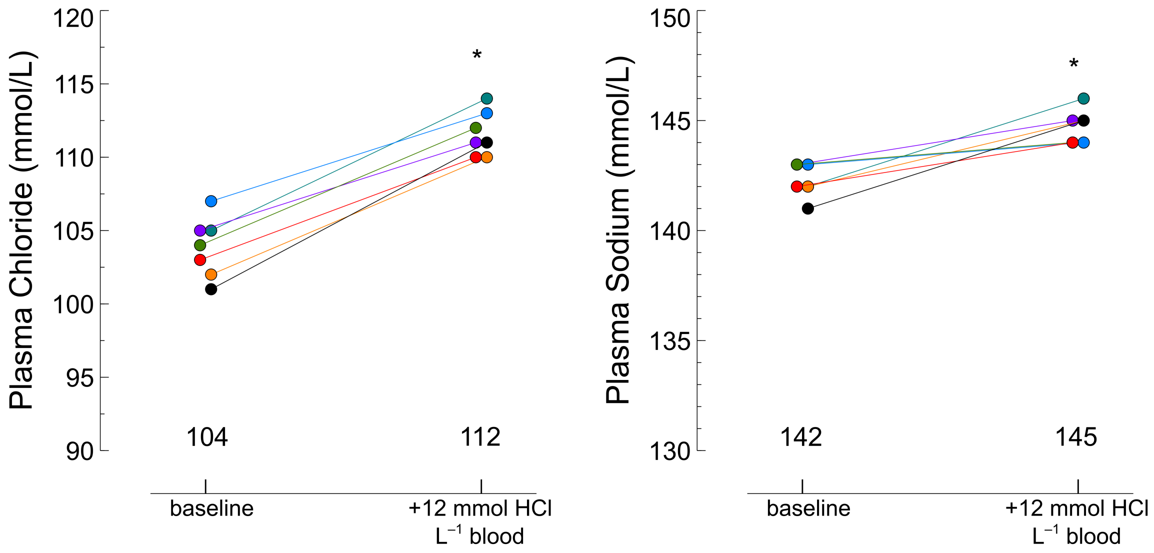


***In vitro* titration of arterial blood with hydrochloric acid**. Arterial blood samples were obtained from a single Sprague-Dawley rat and immediately analyzed without and with the addition of 6 µmol HCl to 500 µL blood (i.e. 20 mmol chloride per L plasma for an hematocrit of 40%). The average hematocrit was 44%. *) $P < 0.05$ compared to baseline.

**Supplemental Table 1.** ***Normal intervals***

| **Parameter** | **Normal interval †** |
|---|---|
| pH | 7.35 – 7.45‡ |
| Actual plasma bicarbonate (mmol/L) | 22 – 27* |
| Partial pressure of $CO_2$ (*p*CO2; kPa) | 4.6 – 6.0 |
| Standard Bicarbonate (mmol/L) | 22 – 27 |
| Standard Base Excess (mmol/L) | -3 – +3 |
| Blood Base Excess (mmol/L) | -3 – +3** |
| Strong Ion Difference Gap (ΔSID; mEq/L) | -3 – +3 |

† For arterial blood. We assumed different levels in central venous blood: about 1 mmol/L higher for SBE, 0.033 lower for pH, 0.6 kPa higher for $pCO_2$ and 5 kPa lower for $pO_2$.

‡ In our experience, values between 7.30 - 7.50 are acceptable in clinical practice.

* assuming a pCO2 of 5.3 kPa (40 mmHg)

** assuming a blood Hb of 145 g/L

### REFERENCES


1. Jorgensen K, Astrup P. Standard bicarbonate, its clinical significance, and a new method for its determination. *Scand J Clin Lab Invest* 1957; **9:** 122-132.
